# A general spectral decomposition of causal influences applied to integrated information

**DOI:** 10.1101/629014

**Authors:** Dror Cohen, Shuntaro Sasai, Naotsugu Tsuchiya, Masafumi Oizumi

## Abstract

Quantifying causal influences between elements of a system remains a central topic in many fields of research. In neuroscience, causal influences among neurons, quantified as integrated information, have been suggested to play a critical role in supporting subjective conscious experience. Recent empirical work has shown that the spectral decomposition of causal influences can reveal frequency-specific influences that are not observed in the time-domain. To date however, a spectral decomposition of integrated information has not been put forward. In this paper, we propose a spectral decomposition of integrated information in linear autoregressive processes. Our proposal is based on a general and flexible framework for deriving the spectral decompositions of causal influences in autoregressive processes. We show that the framework can retrieve the spectral decompositions of other well-known measures such as Granger causality. In simulation, we demonstrate a complex interplay between the spectral decomposition of integrated information and other measures that is not observed in the time-domain. We propose that the spectral decomposition of integrated information will be particularly useful when the underlying frequency-specific causal influences are masked in the time-domain. The proposed method opens the door for empirically investigating the relevance of integrated information to subjective conscious experience in a frequency-specific manner.

**Author summary:** Understanding how different parts of the brain influence each other is fundamental to neuroscience. Integrated information measures overall causal influences in the brain and has been theorized to directly relate to subjective consciousness experience. For example, integrated information is predicted to be high during wakefulness and low during sleep or general anesthesia. At the same time, neural activity is characterized by well-known spectral signatures. For example, there is a prominent increase in low frequency power of neural activity during sleep and general anesthesia. Taking account of the spectral characteristics of neural activity, it is important to separately quantify integrated information at each frequency. In this paper, we propose a method for decomposing integrated information in the frequency domain. The proposed framework is general and can be used to derive the spectral decomposition of other well-known measures such as Granger causality. The spectral decomposition of integrated information we propose will allow empirically investigating the relationship between neural spectral signatures, integrated information and subjective consciousness experience.

## Introduction

Across diverse scientific fields, characterizing the nature of interactions among elements of a system is fundamental to understanding complex, system-level phenomena. In neuroscience, complex interactions among many neurons are thought to give rise to system-level phenomena such as attention [1], learning [2] and even conscious experience [3]. To understand such complex interactions, modern analysis tools often seek to measure the degree of causal influence among nodes or mechanisms of the system. Here, we use the term “causal” to refer to effects across time (e.g. the effect of neuron A at time *t* on neuron B at time *t* + *τ*) [4]. Causal influence are contrasted with equal-time or correlational influences (e.g. the effect of neuron A at time *t* on neuron B at time *t*). A common way to quantify causal influences is by using the technique of transfer entropy [5] or its equivalent under the Gaussian assumption, Granger causality [6–8].

Granger causality has been extensively used in the neurosciences to study causal influences between different brain areas, and how these are modulated by cognitive processes such as attention [4, 9, 10]. In recent years, there has been an increasing interest in investigating causal influences in the frequency domain. To this end, the spectral decomposition of Granger causality developed by Geweke [7, 11, 12] has been instrumental. For example, [13] and [14] used the spectral decomposition of Granger causality to show that feedforward and feedback in the monkey and human visual cortex are mediated by higher and lower frequencies respectively.

The quantification of causal influences is also playing an increasingly important role in the neuroscience of consciousness. For example, a number of studies have shown that transfer entropy from frontal to parietal brain areas is disconnected during unconsciousness as induced by various general anesthetics [15–17]. Recently, we have reported that general anesthesia reduces low-frequency feedback from the center to the periphery of the fly brain, while high-frequency feedforward in the opposite direction remained intact [18].

The increasing prominence of integrated information theory is further emphasizing the importance of investigating causal influences in the brain. Integrated information theory claims that the amount of integrated information - the information a system generates causally as a whole, above and beyond the amount of information generated by its parts independently - is fundamentally related to the subjective conscious experience of that system [3, 19–21].

Quantifying and empirically investigating integrated information, termed Φ, has been the subject of increasing interest. There are currently numerous ways to calculate integrated information [19–26]. In our previous work [22], we demonstrated that a single unified framework can be used to derive integrated information as well as stochastic interaction [27], predictive information [28] and transfer entropy [5] in the time-domain.

In spite of the theoretical and empirical interest in measuring integrated information, there are currently no known spectral decompositions for this quantity. This may be a serious oversight since neural activity is known to display complex frequency-by-frequency behavior, as noted above. Providing a spectral decomposition of integrated information may be particularly important since neural activity in the conscious and unconscious brain can be distinguished by the activity’s spectral characteristics. For example, the prominent increase in low frequency power observed during deep sleep [29–32]. In this paper, we propose a spectral decomposition of integrated information. Our proposal extends our previous work [22] by focusing on autoregressive processes and their representation in the frequency domain. In [22], we defined the time-domain integrated information as the Kullback-Leibler (KL) divergence between the probability distribution in a fully-connected model and that in a disconnected model where causal influences between the elements were removed (Eq. 11 in [22]). Further, we showed that under autoregressive assumptions, integrated information defined in this way was equivalent to the log ratio of the prediction errors in the disconnected and fully connected model (Eq. 23 in [22]). Here, we first generalize this equivalence to autoregressive processes for which an arbitrary number of time steps is considered and any subset of causal influences is removed when forming the disconnected model. Next, using a known result that connects the spectral density matrix to the prediction error [7], we present a general spectral decomposition. Finally, we use this general spectral decomposition to present a spectral decomposition of integrated information. The spectral decomposition is such that the integral over frequencies is equal to the time-domain measure. We demonstrate that by using the general spectral decomposition with respect to different disconnected models, we can (1) derive the spectral decomposition of Granger causality proposed by [7], (2) show that the spectral decomposition of stochastic interaction [27] is closely related to coherence, and derive spectral decompositions for (3) predictive information and (4) instantaneous interaction. We show using simulations that the spectral decompositions of integrated information, Granger causality and stochastic interaction display a complex frequency-by-frequency interplay that is not observed in the time-domain. This demonstrates that the spectral decomposition of integrated information reveals information that is masked in the time-domain.

## Results

In this section, we present our approach for quantifying causal influences and their spectral decompositions and put forward our framework for the spectral decomposition of integrated information.

### Full and disconnected multivariate autoregressive processes

#### Full model

We begin by considering a multivariate, stochastic dynamical system in which the states of the system at times *t* to *t* − *p* are given by ***x***^*t*^, ***x***^*t*−1^, ***x***^*t*−2^, *…* ***x***^*t*−*p*^. The vector ***x***^*t*−*k*^ represents the states of *N* elements at time *t* − *k*,

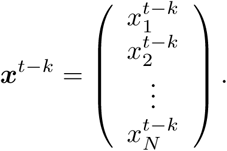

We assume that the system can be represented by the standard pth-order, multivariate linear autoregressive model [33]

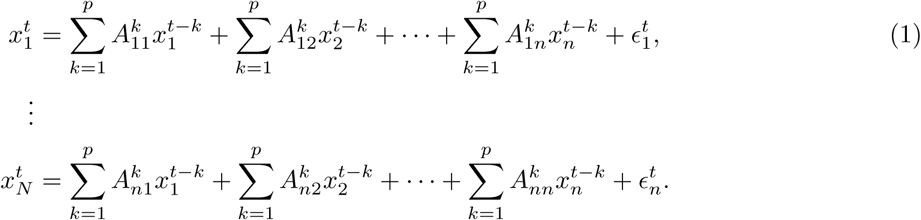

By collecting the autoregressive coefficients into a matrix, the autoregressive model can be compactly written

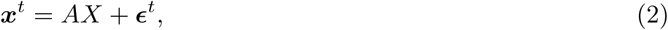

where

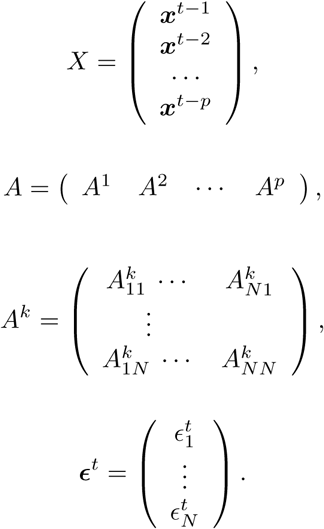

The matrix *A*^*k*^ is called a connectivity matrix and its elements 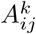 capture how past values of the element *j* influence *k*-step future values of the element *i*. We will refer to these as *causal* influences. 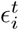 represent normally distributed residuals that are uncorrelated over time. The covariance matrix of the residuals is denoted by Σ(***ϵ***^*t*^) where 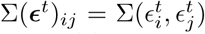. Since the residuals are uncorrelated over time we drop the superscript and use Σ(***ϵ***) and Σ(***ϵ***)_*ij*_. We will refer to Σ(***ϵ***)_*ij*_, *i* ≠ *j*, as *instantaneous* influences since they correspond to equal time correlations.

We use Γ^*k*^ to denote the auto-covariance of ***x***^*t*^ and ***x***^*t*−*k*^,

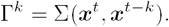

By using Γ^*k*^, the covariance of *X* and ***x***^*t*^ is

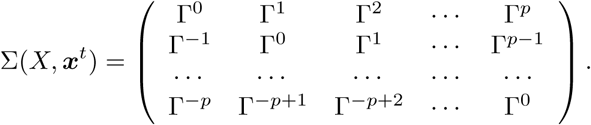

The accuracy of predicting the present state ***x***^*t*^ given the past states *X* is quantified by the determinant of Σ(***ϵ***), |Σ(***ϵ***)|, which is called generalized variance [8]. The smaller |Σ(***ϵ***)| is, the more accurate the prediction is. Given the autocovariance of the system Γ^*k*^, we can find the optimal connectivity matrix *A* that minimizes the generalized variance by differentiating |Σ(***ϵ***)| with respect to all the components of the connectivity matrix, 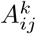[24]. Minimizing |Σ(***ϵ***)| with respect to *A* corresponds to the maximum likelihood estimation [24],

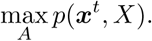

Rewriting the covariance matrix of the residuals using Eq. 2 gives

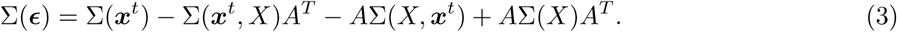

Differentiation of |Σ(***ϵ***)| is calculated as follows

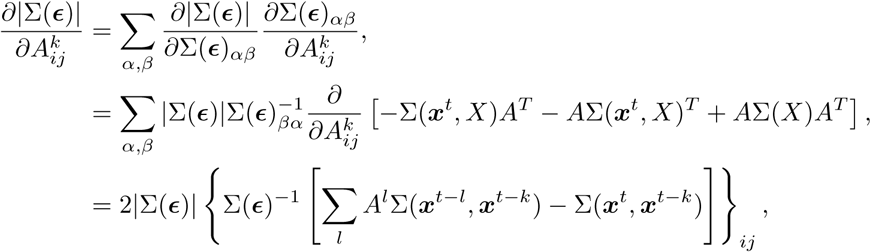

where we used the formula

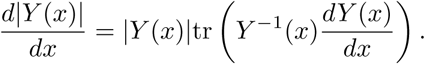

Thus, the optimal *A* that minimizes the generalized variance is obtained by solving the following equations,

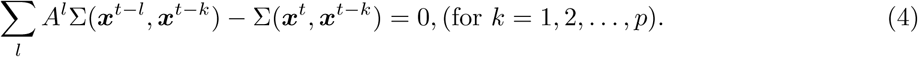

These are the well-known Yule-Walker equations, for which efficient solvers are available [34]. Note that optimizing A with respect to the trace of Σ(*ϵ*) (which is equivalent to minimizing the mean square error of the residuals) also leads to Eq. 4 [8].

We refer to the above model (Eqs. 1 or 2) as the ‘full model’ since we have placed no constraints on the connectivity matrix *A*, i.e., every element *i* in the system can causally affect every other element *j* through a non-zero entry 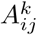 (Figure 1A).

**Figure 1.**
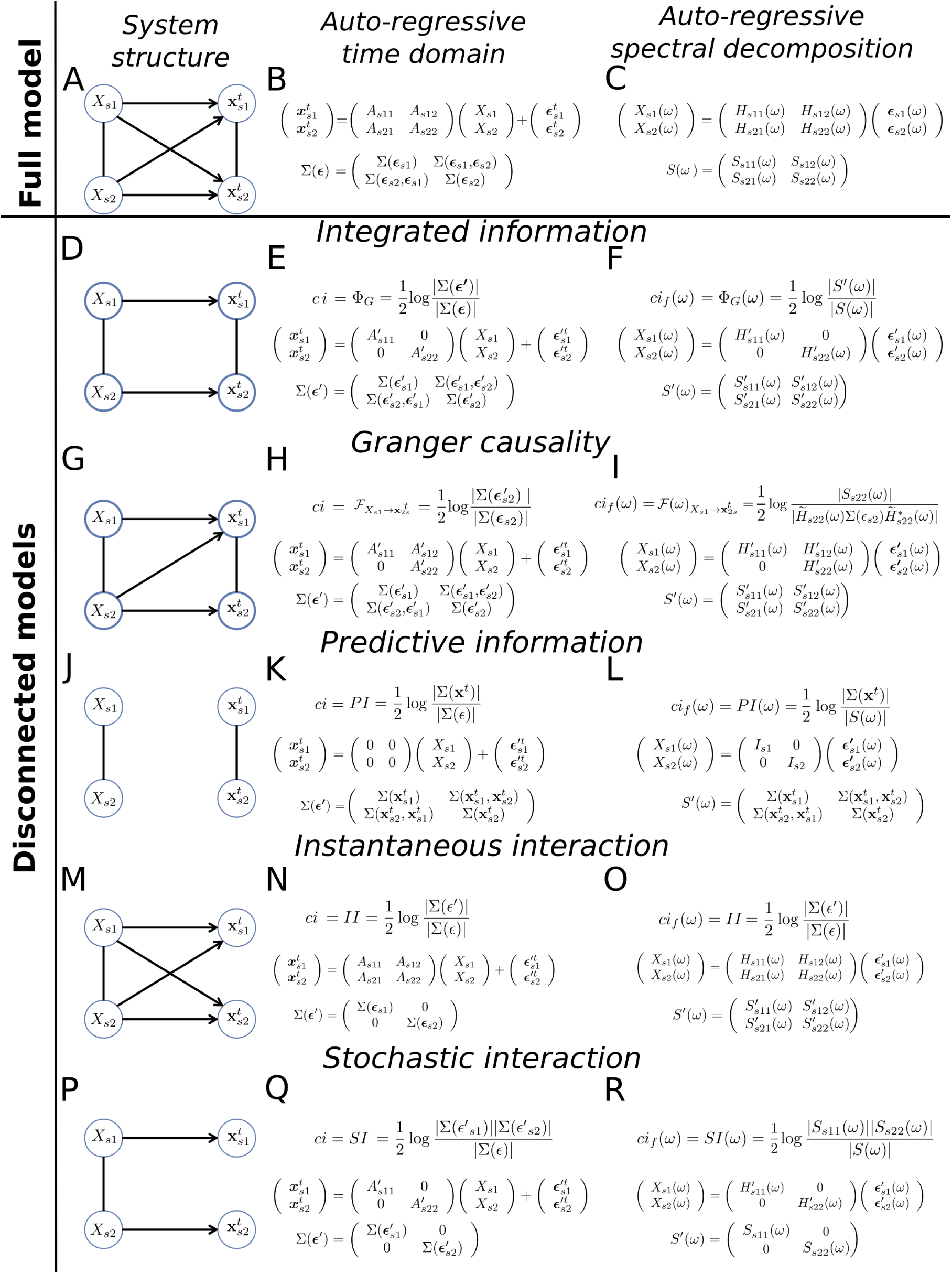
General framework for quantifying causal influences and their spectral decompositions. By comparing the generalized variances and the determinants of spectral density matrices of the full against disconnected models we obtain different measures of causal influences and their spectral decompositions. **A**) Structure of the full model. Horizontal lines indicate influences from the past to the present of each node. Diagonal lines indicate influences from the past of one node to the present of the other (causal influences). The vertical line connecting 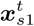 and 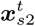indicates instantaneous influences. The vertical line connecting *X*_*s*1_ and *X*_*s*2_ represents the autocovariance function of the system. Note that *X* describes the states of the system in the past k steps, whereas ***x***^*t*^ refers to the “single” time step at the present. **B**) The autoregressive model for the system in **A** is represented by the connectivity matrix *A* and the covariance of the residuals Σ(***ϵ***). **C**) In the frequency domain the autoregressive model is represented by the transfer function *H*(*ω*) and spectral density matrix *S*(*ω*). **D**) The disconnected model used to define integrated information contains no causal influences. **E**) The corresponding autoregressive model has a block-diagonal connectivity matrix *A′*. Σ(***ϵ****′*) is not constrained. We quantify causal influences *ci* using the log ratio of the generalized variance of the full and disconnected models, here leading to integrated information Φ_*G*_. **F**) The spectral representation of the model has a block-diagonal transfer function *H′* (*ω*). The spectral decomposition of the causal influences (*ci*_*f*_) is defined as the log ratio of the determinants of the spectral density matrices (see text for details), leading to the spectral decomposition of integrated information Φ_*G*_ (*ω*). By considering different disconnected models (**G, J, M** and **P**) we obtain the time-domain autoregressive models corresponding to Granger causality (**H**), predictive information (**K**), instantaneous interaction (**N**) and stochastic interaction (**Q**), as well as the corresponding frequency-domain representations used in their spectral decompositions (**I, L, O** and **R**, see text for details).

#### Disconnected models

Next, we consider a “disconnected” model in which some influences are “cut”. We cut influences by forcing some of the causal influences to be zero. We denote the disconnected connectivity matrix and corresponding residuals covariance matrix by *A′* and Σ(*∊*′), respectively. The auto-regressive model of the disconnected model can be written analogously to Eq. 2 as

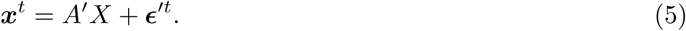

In general, we can cut any combination of causal influences. We introduce the set Λ which includes all the combinations of the influences from node (or set of nodes) *m* to node (or set of nodes) *n* that are cut. When cutting influences across all lags, the constraints are generally expressed as

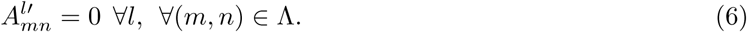

As before, we can minimize the prediction error of the disconnected model by minimizing the generalized variance |Σ(***ϵ****′*) | with respect to the connectivity matrix *A′*. First, the entries of the covariance matrix of the residuals of the disconnected model are given by

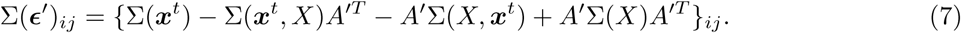

We find the optimal connectivity matrix, which minimizes the prediction error, by differentiating |Σ(***ϵ****′*)| with respect to 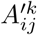,

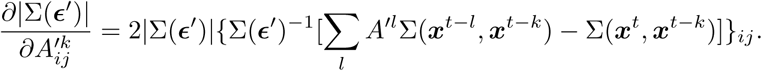

Thus, the optimal *A′* that minimizes the generalized variance satisfies the following

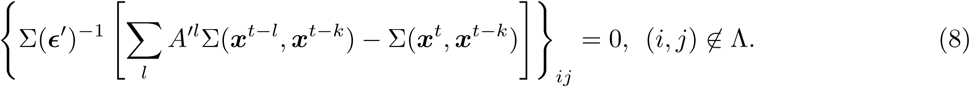

The optimal connectivity matrix *A′* is obtained by solving Eqs. 7 and 8 under the constraints given by Eq. 6.

### General framework for spectral decomposition of causal influences

#### Quantifying causal influences in the time domain

In our previous work [22], we proposed to quantify causal influences by evaluating the Kullback-Leibler (KL) divergence between the probability distributions of the full and disconnected models. In the multivariate autoregressive model (Eqs. 2 and 5), the probability distributions of the full model *p* and the disconnected model *q* are given by

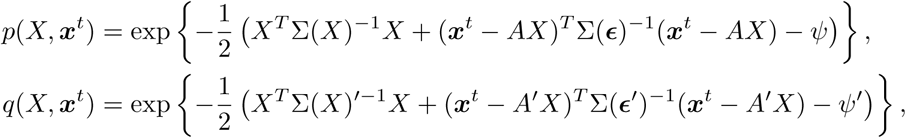

where *ψ* and *ψ ′* are normalization factors which ensure that the integral of the probability distributions evaluates to 1. The disconnections in the model mean that *q* is constrained in some way. For example, if we cut the causal influences from node *n* to node *m*, the constraints we impose on 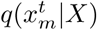 are given by [22]

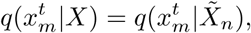

where 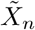 is the complement of *X*_*n*_ in the whole system *X*. This constraint means that there is no direct influence from the node *n* to node *m* given the states of the other elements 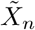 being fixed. In terms of the connectivity matrix *A′*, this constraint is expressed as (see Supplementary information)

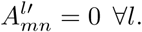

The causal influences (*ci*) are quantified by minimizing the KL divergence between *p* and *q* under the constraints of the disconnected model,

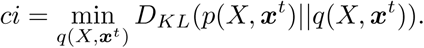

Minimizing the KL divergence corresponds to finding the best approximation of the full model *p* using the constrained distribution *q*. We can find the minimizer of the KL divergence by differentiation with respect to the parameters of the disconnected model Σ(*X*)^*′*−1^, Σ(***ϵ****′*)^−1^, *A′* [22],

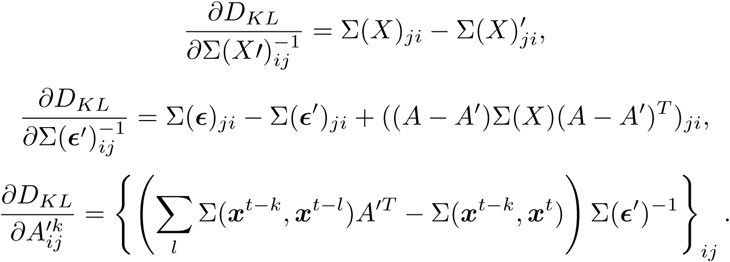

By setting the derivatives of the KL divergence to 0, we have the following equations that give the minimizer of the KL divergence

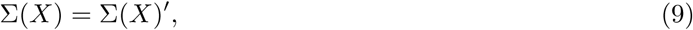

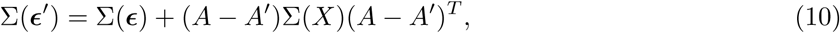

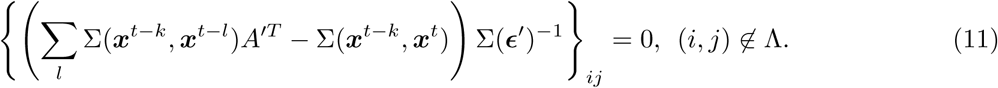

By substituting Eq. 3 into Eq. 10 and using Σ(***x***^*t*^, *X*) = *A*Σ(*X*), we can see that Eq. 10 is equivalent to Eq. 7. Also, Eq. 11 is simply a transposed version of Eq. 8. This means that minimizing the KL divergence is equivalent to minimizing the prediction error |Σ(***ϵ****′*) |. By substituting Eqs. 9 and 10 into the KL divergence, the minimized KL divergence is expressed as

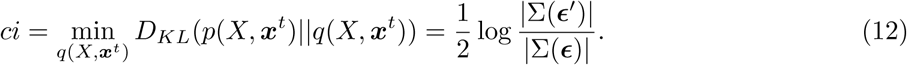

Σ(***ϵ****′*) is obtained by solving Eqs. 10 and 11 under the constraints (Eq. 6). In terms of prediction error, we can interpret the proposed measure as the log difference of the minimized prediction error in the full and disconnected models. Note that the expression in Eq. 12 holds for any constraints given by Eq. 6. In [22] we showed that integrated information, Granger causality, and predictive information can all be derived by using Eq. 12 with respect to different constraints (i.e. different disconnected models).

In the next section, we derive a general spectral decomposition of Eq. 12 by using a key relationship that connects the generalized variance |Σ(***ϵ***) | to the spectral density matrix *S*(*ω*). We will use this general spectral decomposition to define a spectral decomposition of integrated information.

#### Spectral decomposition of causal influences

To derive the spectral decompositions of causal influences, we first express the autoregressive process in the frequency domain (see [7, 11, 12, 34, 35] for further details). By introducing the lag operator *A*(*L*) we rewrite Eq. 2 as

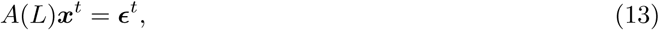

where

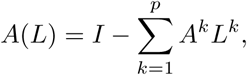

*I* is the identity matrix and *L* is the lag operator,

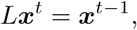

Fourier transforming both sides of Eq. 13 leads to

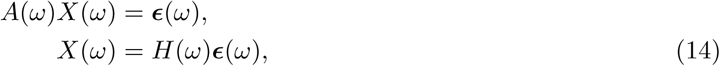

where

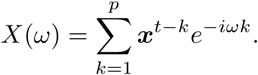

*A*(*ω*) is an *N* × *N* matrix with entry at row *l* and column *m* given by

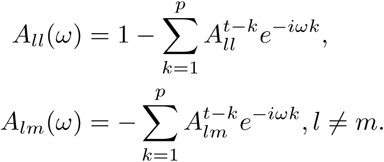

The transfer function is defined as

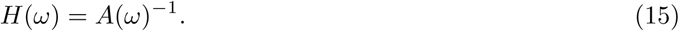

The Fourier transform of the residuals is

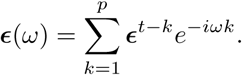

We obtain the spectral density matrix *S*(*ω*) by multiplying by the complex conjugate of both sides of Eq. 14,

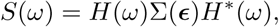

where ***ϵ*** (*ω*) ***ϵ*** ^*\**^(*ω*) = Σ(***ϵ***) because ***ϵ***^*t*^ are Gaussian random variables that are uncorrelated over time. Under the condition that the autoregressive model (Eq. 2) has a stationary solution (which is equivalent to the condition that all roots of |*A*(*L*)| lie outside the unit circle), the following holds [7]

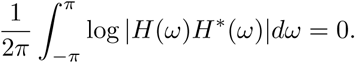

Thus, we have the relation

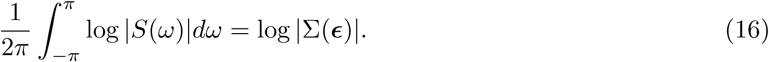

This important relationship connects the frequency-by-frequency characteristics of the system, as described by |*S*(*ω*)|, to the overall prediction error, as quantified by the generalized variance |Σ(***ϵ***)|, in the time-domain. This relationship forms the basis of our proposed spectral decomposition of integrated information.

Following analogous steps for the disconnected autoregressive model (Eq. 5), we have

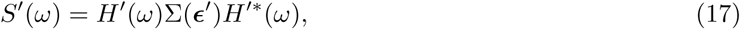

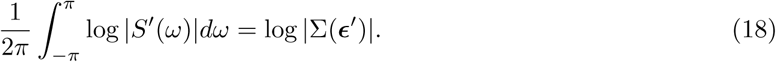

From Eqs. 16 and 18, we can recover the time-domain measure (Eq. 12) through integration over frequencies

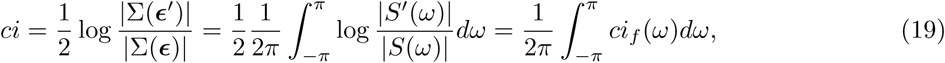

where *ci*_*f*_ (*ω*) is defined as

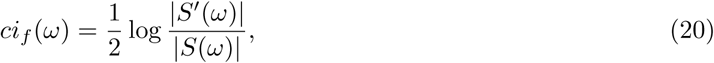

*ci*_*f*_ (*ω*) is interpreted as the spectral decomposition of causal influences. The lower subscript *f* represents “frequency”. Note that *ci*_*f*_ (*ω*) is represented by the log ratio of the spectral density matrices of the full and disconnected models, which is analogous to *ci* in the time-domain being represented by the log ratio of the generalized variances of the full and disconnected models.

As we will show next, we can use this framework to define a spectral decomposition for integrated information. We will also apply this framework to derive the spectral decomposition of Granger causality, predictive information, instantaneous interaction and stochastic interaction.

### Spectral decomposition of integrated information

Integrated information quantifies total causal influences between subsystems. In order to quantify integrated information, we first need to partition the system into subsystems of interest. Here we consider a bipartition; grouping the elements into two subsystems, 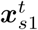 and 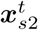. We emphasize, however, that our approach is directly applicable to *any* partition scheme that can be defined by cutting a subset of the causal influences. The full autoregressive model (Eq. 2) with respect to this grouping can be expressed as (Figure 1B),

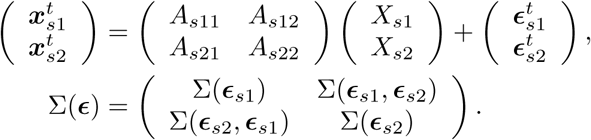

The representation of the autoregressive model in the frequency domain is given by (Figure 1C),

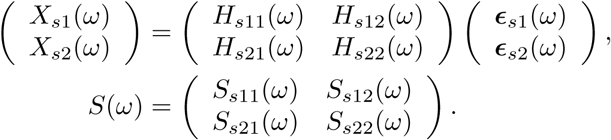

To quantify integrated information, termed Φ_*G*_, we consider a disconnected model in which all causal influences between the subsystems are cut (Figure 1D). The constraints on the probability distribution are [22]

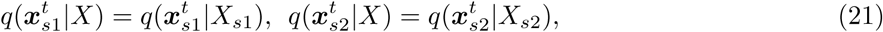

which are equivalent to the following constraint on *A ′* (see Supplementary information)

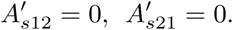

The corresponding disconnected model thus has a block-diagonal connectivity matrix (Figure 1E),

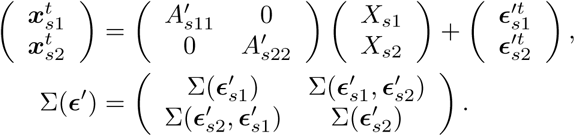

As we showed, the minimized KL divergence between the full and disconnected model under the constraints (Eq. 21) corresponds to the log ratio of the generalized variances (Eq. 12),

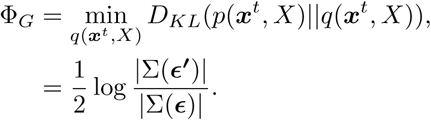

The disconnected connectivity matrix *A ′* and covariance of residuals Σ(***ϵ′***) are estimated by iteratively solving Eqs. 7 and 8.

The corresponding representation of the autoregressive model in the frequency domain is

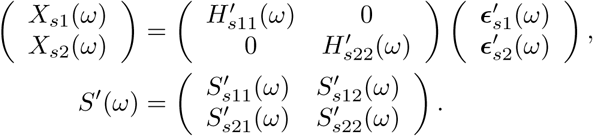

where 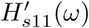 and 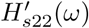 are estimated using *A′*(*ω*) instead of *A*(*ω*) (Eq. 15). The spectral density matrix of the disconnected model *S′*(*ω*) is obtained using the estimated Σ(***ϵ ′***) and *H′*(*ω*) (Eq. 17).

Our proposed spectral decomposition of Φ_*G*_ (Eq. 20) is

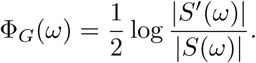

### Spectral decomposition of other measures of causal influence

Next, we apply the same framework to derive the spectral decompositions of Granger causality, predictive information, instantaneous interaction and stochastic interaction. The derivations of Granger causality and predictive information are straightforward since we can directly use the equivalence between the minimized KL and the log ratio of generalized variances we derived (Eq. 12). The derivation of instantaneous interaction and stochastic interaction involves constraints on the covariance matrix of the residuals, and thus we cannot directly use the equivalence between the minimized KL and the log ratio of generalized variances (Eq. 12). We address this limitation in the “Spectral decomposition of instantaneous interaction and stochastic interaction” section.

#### Granger causality

Granger causality measures how much past values of one variable improve predictions of future values of another variable. To quantify this in our framework, we consider a disconnected model in which all causal influences in one direction are cut. For example, to evaluate Granger causality from *X*_*s*1_ to 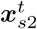, we cut all causal influences from *X*_*s*1_ to 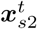 (Figure 1G). The constraint on *q* is given by

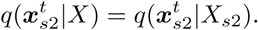

From the autoregressive perspective, the constraint on *q* is equivalent to forcing the bottom, off-diagonal sub-matrix of *A′* to zero (see Supplementary information) (Figure 1H). The disconnected model is

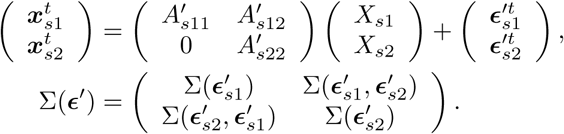

The minimized KL divergence under this constraint is equivalent to transfer entropy from *X*_*s*1_ to 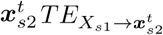

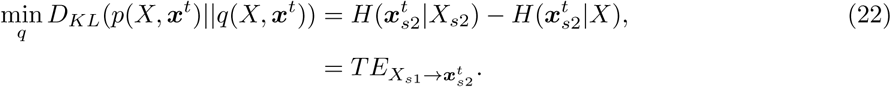

Under the Gaussian assumption, transfer entropy equals Granger causality [36], denoted 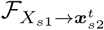

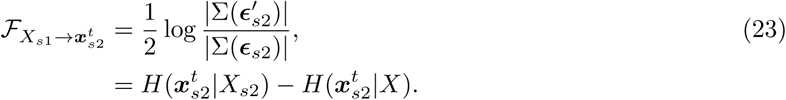

Eq. 23 is the standard definition of Granger causality, where the generalized variance in one of the subsystems is compared between the full model and the disconnected model. From Eq. 12, this equals the log ratio of the generalized variances in the whole system;

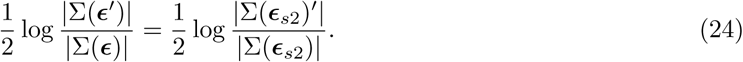

The frequency domain description of the disconnected model is (Figure 1I)

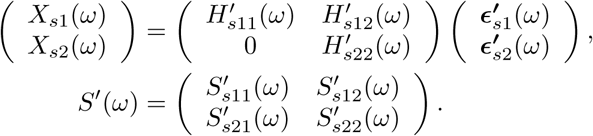

Using our proposed spectral decomposition (Eq. 20), the spectral decomposition of Granger causality is given by

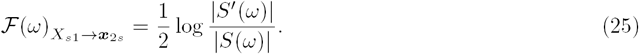

Previously, the spectral decomposition of Granger causality from *X*_*s*1_ to 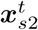 was defined as [7]

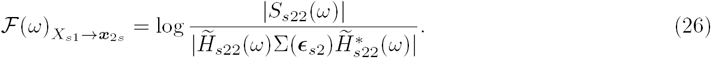

where 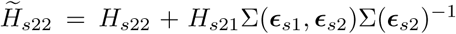. At first sight the spectral decomposition of Granger causality as derived by our framework (Eq. 25) appears different from that suggested by Geweke (Eq. 26). However, we can show that these are indeed equal (see Supplementary information),

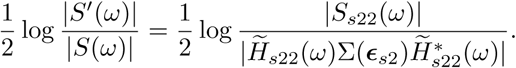

The difference between the conventional approach taken by Geweke [7] and our framework is that in the former the variance of the *subsystems* is compared (rhs of Eq. 24), whereas in our approach the variance of the *entire* system is compared (lhs of Eq. 24). Focusing on the entire system is the more general approach as this also allows deriving other measures such as integrated information and predictive information in a systematic way (see the following sections), and is also directly extensible to partitions other than bipartitions. Furthermore, the spectral decomposition in our framework is easily derived by using a simple equation (Eq. 19), while the original derivation of the spectral decomposition given by Geweke ([7]) is complicated.

#### Predictive information

Next we consider a disconnected model in which all causal influences are cut (Figure 1J). The constraint on *q* is

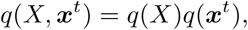

where *q*(*X*) and *q*(***x***^*t*^) represent the marginal distributions of *X* and ***x***^*t*^ under *q*(*X*, ***x***^*t*^). The constraint on the autoregressive model is *A*′ = 0 (see Supplementary information), leading to the disconnected model (Figure 1K)

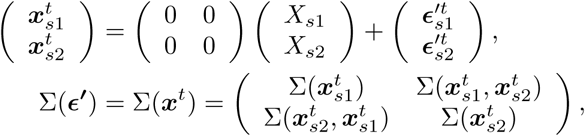

where the last equality follows from *A*′ = 0 (by substitution in Eq. 7).

The minimized KL reduces to predictive information, termed *PI*,

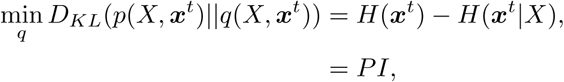

which is equivalent to mutual information between the past and the present, *MI*(***x***^*t*^; *X*)). We find

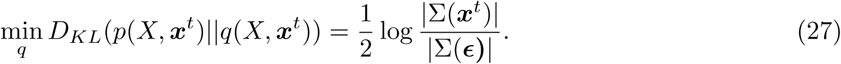

In the frequency domain, we have *H*′(*ω*) = *I*, from using *A*′ = 0 and the definition of the transfer function (Eq. 15). Thus, the frequency domain representation of the disconnected model is (Figure 1L)

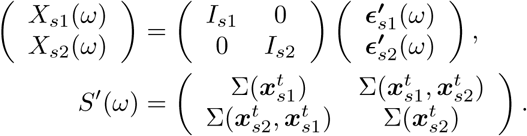

The form of *S*′(*ω*) follows from

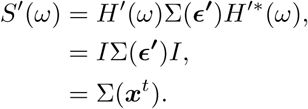

Thus, the spectral decomposition of predictive information is given by

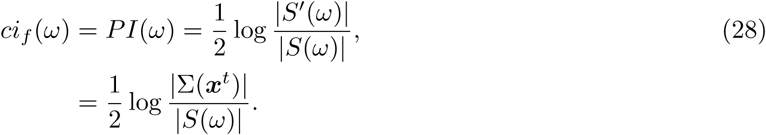

Note that log |Σ(***x***^*t*^)| is constant across frequencies and that for a univariate process log |*S*(*ω*)| corresponds to the (log of the) power spectrum. Intuitively, deviations from log |Σ(***x***^*t*^)| at a particular frequency suggest that the past exerts some influence on the future at that particular frequency. Note that this quantity can become negative if |*S*(*ω*) | *>|* Σ(***x***^*t*^)| (see Relationship between the measures in the frequency domain, Figure S1).

### Spectral decomposition of instantaneous interaction and stochastic interaction

In this section we investigate how our framework relates to two other well known measures-instantaneous interaction ([7, 11, 12]) and stochastic interaction ([37]). For the measures of causal influences we considered in the previous section the constraints were imposed only on the connectivity matrix *A*. Instantaneous interaction and stochastic interaction involve constraints on the structure of the residual covariance matrix Σ(ϵ). Because the equivalence between the minimized KL divergence and the log ratio of generalized variances (Eq. 12) is derived under constraints imposed only on the causal influences, we cannot use this equivalence to derive instantaneous interaction and stochastic interaction. However, as we show below, both measures can also be expressed as log ratios of generalized variances through the consideration of specific disconnected models.

#### Instantaneous interaction

Quantifying instantaneous interaction corresponds to quantifying the contribution of equal time influences (Figure 1M). This measure was not derived in [22] so we derive it here. The constraint on the disconnected model is

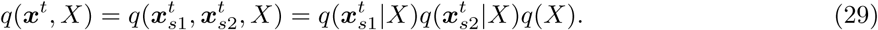

The constraint does not involve the connectivity matrix but requires that the off-diagonal elements of Σ(*ϵ*′) are 0 (see Supplementary information), leading to the following disconnected model (Figure 1N),

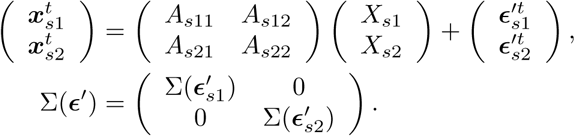

The minimized KL divergence equals instantaneous interaction denoted *II* (see Supplementary information for derivation)

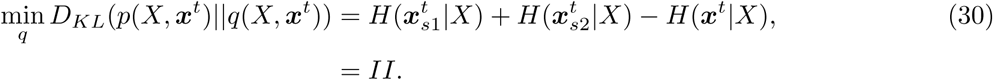

Under the Gaussian assumption, we find

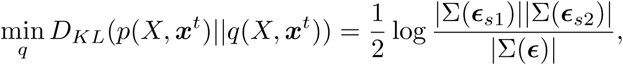

In the Granger causality literature this quantity is known as “instantaneous causality”, denoted 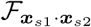 [7, 11, 12, 35]. We prefer the term instantaneous interaction since in our view the term instantaneous *causality* is misleading.

The log ratio of the generalized variances is

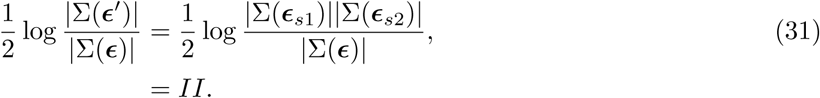

Thus, we see that instantaneous interaction can be expressed as the log ratio of generalized variances in the whole and disconnected model, and so the general spectral decomposition (Eq. 19) can be applied. We emphasize that the diagonal terms in Σ(*ϵ*′) are *identical* to the diagonal terms in Σ(*ϵ*), the difference is in the off-diagonal terms which are forced to zero.

The frequency domain representation of the disconnected model is (Figure 1O)

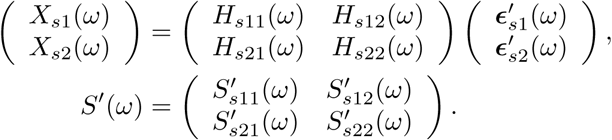

For the spectral decomposition we have

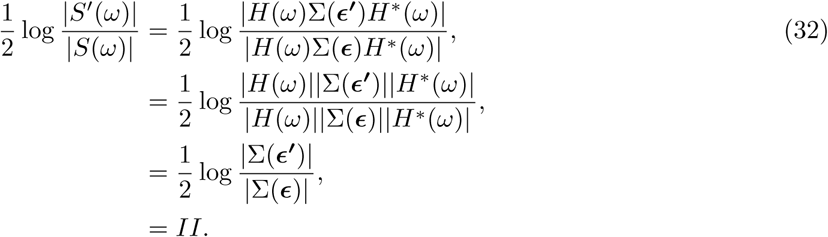

We thus see that the spectral decomposition of instantaneous interactions equals time-domain instantaneous interaction at every frequency. This is unsurprising since the residuals, which are the source of the instantaneous interactions, are assumed to be white. However, we note that the spectral decomposition we derived here is in fact different from that suggested in [11]

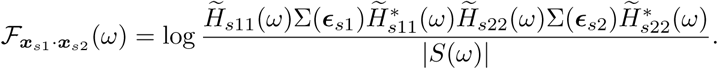

Unlike our derivation of instantaneous interaction (Eq. 32), this measure varies on a frequency-by-frequency basis. However, others have recognized that this quantity lacks theoretical validity, and can become negative in certain situations [35]. In contrast, the measure of instantaneous interaction we systematically derived is always positive and readily interpretable [7].

### Stochastic interaction

Finally, we consider a disconnected model in which both the causal and instantaneous influences between the subsystems are cut (Figure 1P). The constraint on *q* is given by

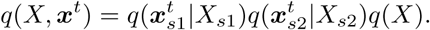

The constraint amounts to a diagonal structure for *A*′ and Σ(*ϵ***′**). The corresponding disconnected model is

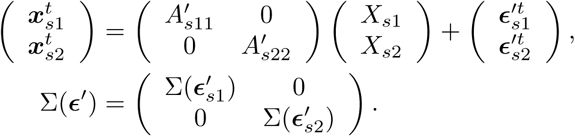

where 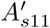 and 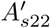 are as estimated when assessing Granger causality from *X*_*s*2_ to 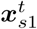 and from *X*_*s*1_to 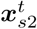, respectively. Intuitively, this disconnected model is obtained by separately fitting autoregressive models to 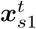 and 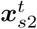 and then simply putting these together to form a ‘complete’ model for 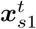 and 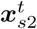.

The minimized KL divergence equals stochastic interaction, denoted *SI*

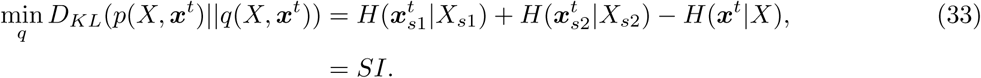

Under the Gaussian assumption this reduces to

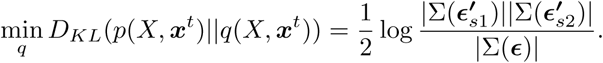

In the Granger causality literature, this quantity is known as “total interdependence”, denoted 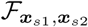, and quantifies overall linear influences [7,11]. Note that 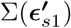 and 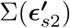 are the same as those estimated when assessing Granger causality from *X*_*s*2_ to ***x***_*s*1_ and from *X*_*s*1_ to 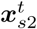, respectively.

We can easily confirm that stochastic interaction can be also written as the log ratio of the determinants of the generalized variances in the whole system

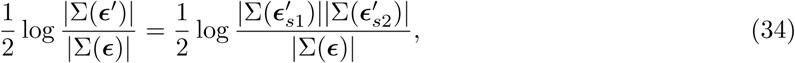

and thus, the general spectral decomposition (Eq. 19) can be applied.

The frequency domain representation of the disconnected model is

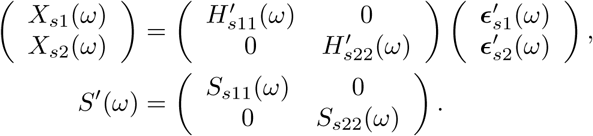

To clarify the reason for the structure of the spectral density matrix, we highlight two points. First, the block diagonal structure of *A*′ and Σ(*ϵ*′) implies a block diagonal structure for *S*′(*ω*), which follows from the definition of *S*′ (*ω*) (Eq. 17). Second, the spectral density matrix of the subprocesses for ***x***_*s*1_ and ***x***_*s*2_ is the same as the corresponding sub-matrices of the full spectral density matrix [7, 38]. That is, we have that 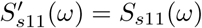 and 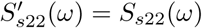.

Finally, the ratio of the determinant of the spectral density matrices is

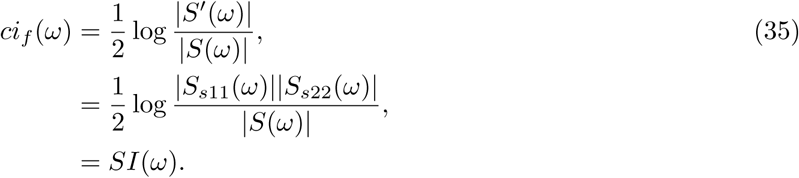

This spectral decomposition is identical to the one obtained for total interdependence 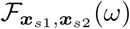 [11]

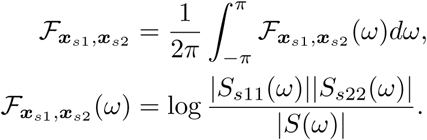

Note that 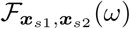 is related to block-coherence *C*(*ω*) [12, 39, 40]

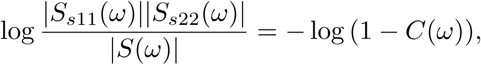

where

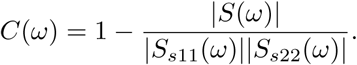

Block-coherence reduces to ordinary coherence in the bivariate case and to multiple-coherence when one of the variables is univariate [39, 40]. We emphasize that the spectral decomposition of total interdependence and its relation to coherence are recognized [7, 11, 35, 39]. What we show here is that (1) this quantity is equivalent to stochastic interaction under the Gaussian assumption and (2) that its spectral decomposition, which is closely related to coherence, can be derived through our general framework.

### Relationship between the measures in the time-domain

In the theoretical section of the paper, we presented a general framework for deriving spectral decompositions and used this framework to present a spectral decomposition of integrated information and other measures. In the time-domain, the measures are analytically related. First, from [22] we have the following relations between integrated information (Φ_*G*_), transfer entropies 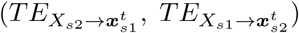, stochastic interaction (*SI*) and predictive information (*PI*)

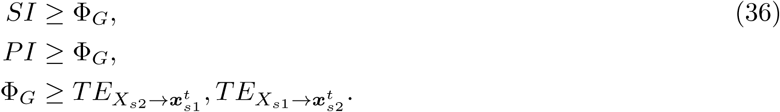

These inequalities reflect that the more influences are cut, the greater the information loss is (Figure 2A).

**Figure 2.**
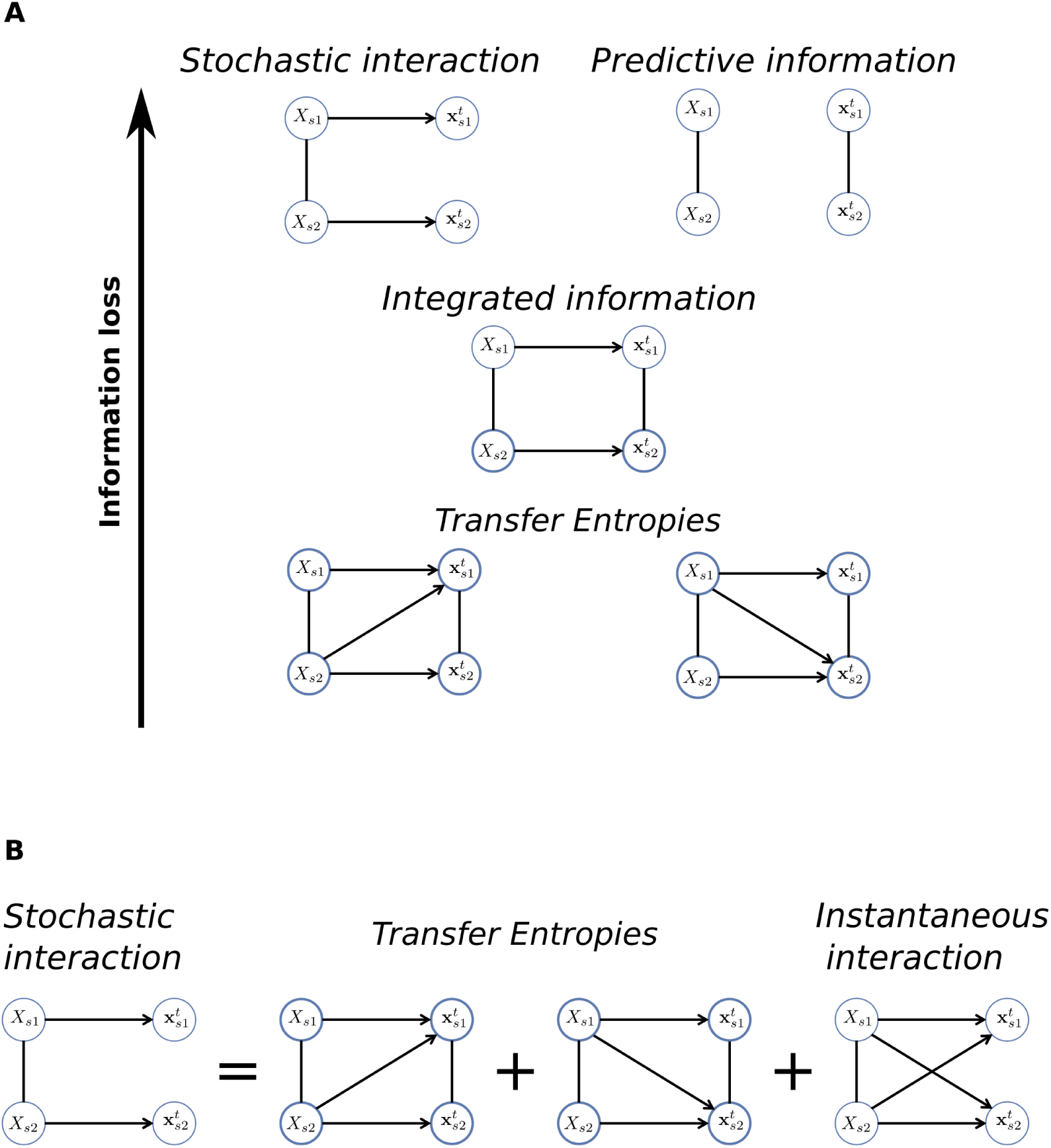
Analytic relationships between the measures in the time-domain. **A**) Schematic demonstrating that the more influences are cut the greater the information loss is (Eq. 36). **B**) Schematic demonstrating that stochastic interaction equals the sum of transfer entropies and instantaneous interaction (Eq. 37).

We also have that the sum of our derived instantaneous interaction (*II*, Eq. 30) and transfer entropies in both directions (Eq. 22) equals stochastic interaction (Eq. 33)

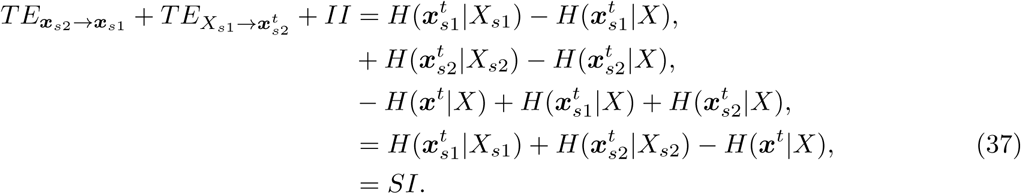

This equality demonstrates that the information loss when cutting both causal and instantaneous influences (stochastic interaction) equals the sum of information losses when cutting the causal (transfer entropies) and instantaneous (instantaneous interaction) influences separately (Figure 2B). Note that a version of this decomposition under the Gaussian assumption is recognized in the Granger causality literature [7, 11]. The derivation above (Eq. 37) shows that the equality holds in the general case.

The intuitive relationships above (Eqs. 36 and 37) help in the interpretation of the measures as meaningful measures of information loss. On the other hand, there are currently no known relationships between the spectral decompositions of these measures. In the following section, we investigate the relationship between the spectral decompositions of the measures using simulations.

### Relationship between the measures in the frequency domain

We used simulations to investigate the relationships between the spectral decompositions of the measures on a frequency-by-frequency basis. We present results from simulations of simple, bivariate autoregressive processes in which the measures and their spectral decompositions can be calculated in a straightforward manner (see Methods).

#### Integrated information and Granger causality are equivalent for an instantaneously-connected and a unidirectionally-connected system

First, we investigated a system with instantaneous but no causal influences between the variables (Figure 3A):

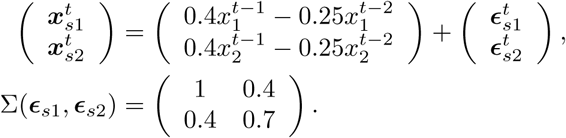

**Figure 3.**
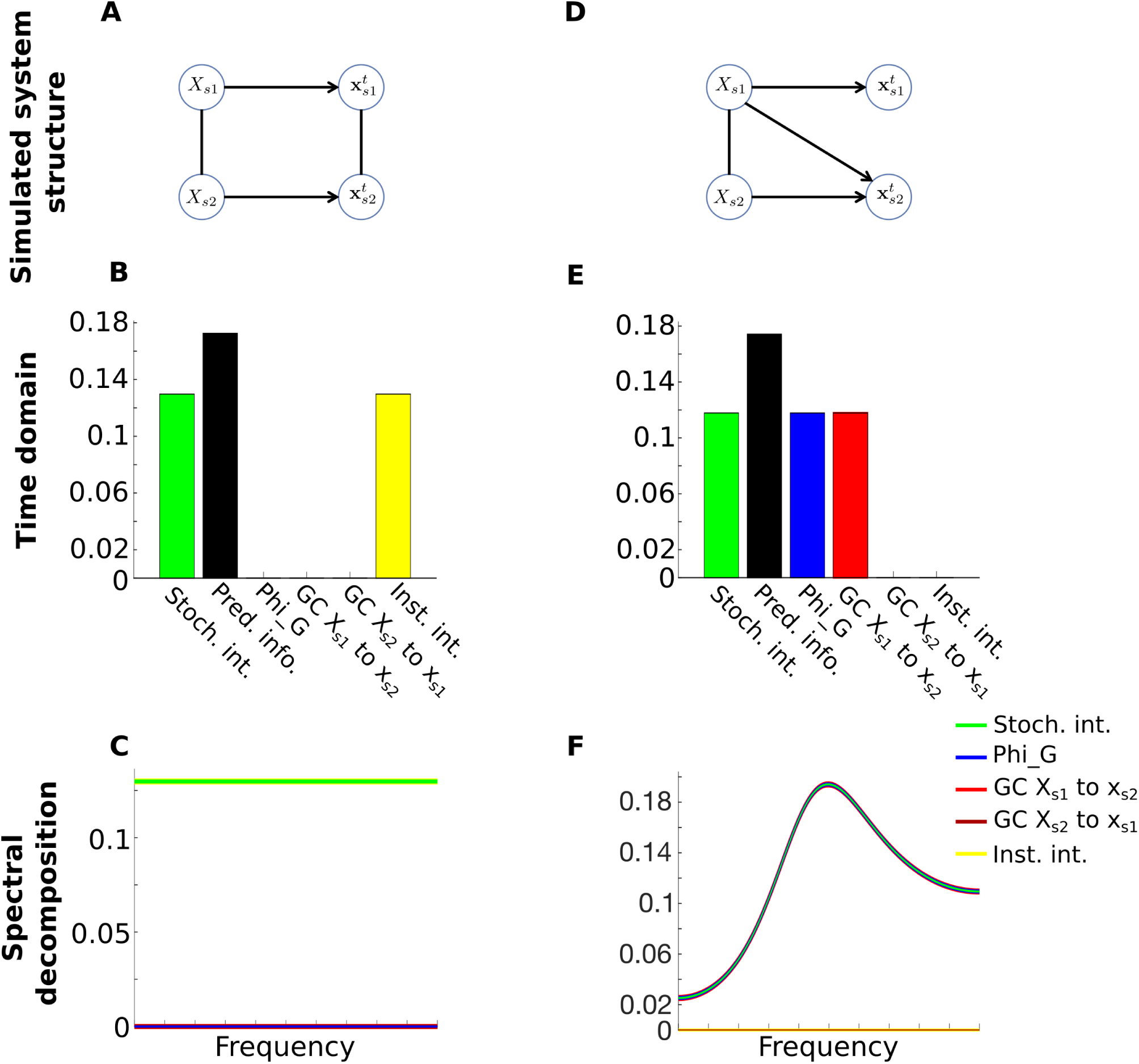
The time-domain relationships (Eqs. 36 and 37) hold on a frequency-by-frequency basis for an instantaneously-connected (**A** - **C**) and a unidirectionally-connected (**D** - **F**) system. **A**) Structure of the instantaneously-connected system. **B**) The measures in the time-domain. Integrated information (blue) = Granger causality from *X*_*s*1_ to **x**_*s*2_ (red) = Granger causality from *X*_*s*2_ to **x**_*s*1_(maroon) = 0. Stochastic interaction (green) = instantaneous interaction (yellow) = 0.130. Predictive information (black) = 0.173. **C**) The spectral decompositions of the measures parallel the time-domain. Integrated information and Granger causality in both directions overlap at *y* = 0 (blue, red and maroon lines overlap). Stochastic interaction and instantaneous interaction overlap at *y* = 0.130. **D**) Structure of the unidirectionally-connected system. **E**) The measures in the time-domain. Integrated information = Granger causality from *X*_*s*1_ to **x**_*s*2_ = stochastic interaction = 0.118. Instantaneous interaction = Granger causality from *X*_*s*2_ to **x**_*s*1_ = 0. Predictive information = 0.174. **F**) The spectral decompositions parallel the time-domain. The spectral decomposition of integrated information, Granger causality from *X*_*s*1_ to **x**_*s*2_ and stochastic interaction (blue, green and red lines) overlap. The spectral decomposition of Granger causality from *X*_*s*2_ to **x**_*s*1_ and instantaneous interaction overlap at *y* = 0. For the spectral decomposition of predictive information for the two systems see Figure S1A and B (see text for details).

The absence of any causal influences means that time-domain integrated information and Granger causality (in either direction) are zero for this system. The system contains self-influences so predictive information is above zero. As per the decomposition of stochastic interaction to the sum of Granger causalities and instantaneous interaction (Eq. 37), we find that stochastic interaction equals instantaneous interaction (Figure 3B).

The spectral decompositions for this system parallel the time-domain. The spectral decompositions of integrated information and Granger causality are zero at every frequency. The spectral decomposition of stochastic interaction and instantaneous interaction equal the time-domain value at every frequency (Figure 3C). In contrast, the spectral decomposition of predictive information has considerably greater magnitude, varies by frequency and attains negative values at some frequencies (Figure S1A). We return to this observation in the Discussion. This simulation demonstrates that, with the exception of predictive information, the time-domain relationships (Eqs. 36 and 37) hold on a frequency-by-frequency basis for this system.

Next, we investigated a system with a unidirectional influence from *X*_*s*1_ to **x**_*s*2_, but no instantaneous influences (Figure 3D):

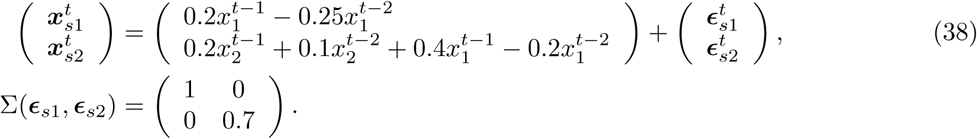

For this system integrated information equals Granger causality from *X*_*s*1_ to **x**_*s*2_, as both quantify the strength of the single causal influence. Due to the absence of any causal influences from *X*_*s*2_ to **x**_*s*1_ or any instantaneous influences, Granger causality from *X*_*s*2_ to **x**_*s*1_ and instantaneous interaction are both zero. Following the time domain relations (Eq. 37), we find that stochastic interaction equals Granger causality from *X*_*s*1_ to **x**_*s*2_ (and thus also equals integrated information). Predictive information is higher than all other measures, quantifying the extent of self-as well as causal-influences (Figure 3E).

For this system, we also find that the spectral decompositions of the measures parallel the time-domain. That is, the equivalence between integrated information, Granger causality from *X*_*s*1_ to **x**_*s*2_ and stochastic interaction in the time-domain is preserved on a frequency-by-frequency basis. Granger causality from *X*_*s*2_ to **x**_*s*1_ and instantaneous interaction equal zero at every frequency. We again observe that the spectral decomposition of predictive information attains negative values at some frequencies (Figure S1B). Thus, we find that similarly to the instantaneously-connected system, the time-domain relationships (Eqs. 36 and 37) hold on a frequency-by-frequency basis for this system, except for predictive information.

#### Integrated information and Granger causality dissociate for a unidirectionally- and bidirectionally-connected system with instantaneous influences

We next investigated the effect of adding instantaneous influences to the unidirectionally-connected system in Figure 3D. We kept the same connectivity matrix *A* (Eq. 38) but added a strong instantaneous influence (Figure 4A) by setting

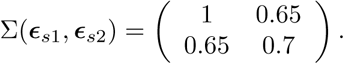

**Figure 4.**
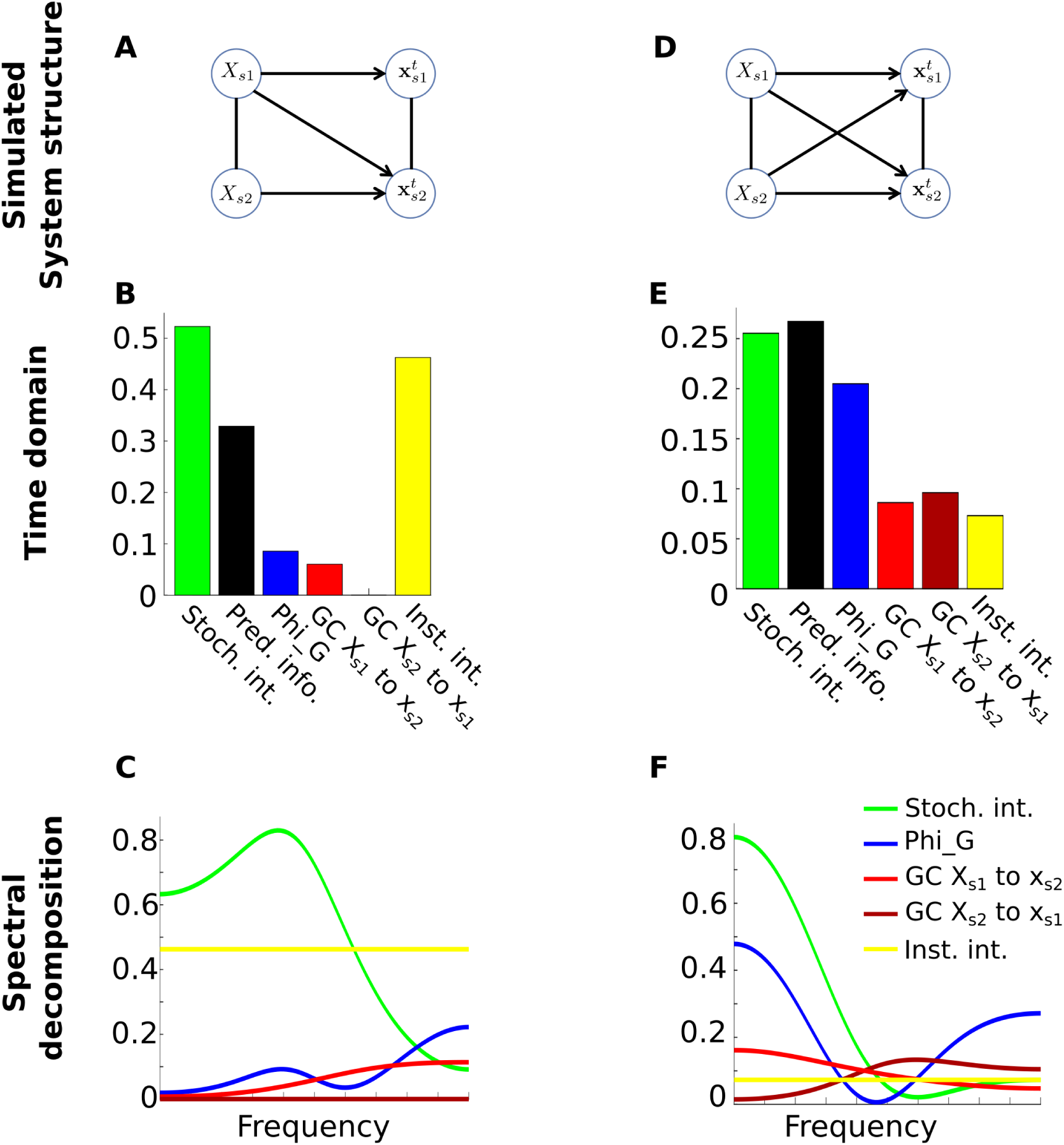
The time-domain relationships (Eqs. 36 and 37) do not hold for a unidrectionally-(**A** – **C**) or a bidirectionally-(**D** – **F**) connected system with an instantaneous influence. **A**) System with a unidirectional influence from *X*_*s*1_ to **x**_*s*2_ and an instantaneous influence between **x**_*s*1_ and **x**_*s*2_. **B**) The measures in the time-domain. Stochastic interaction = 0.523, predictive information = 0.329, integrated information = 0.085, Granger causality from *X*_*s*1_ to *X*_*s*2_ = 0.06, Granger causality from *X*_*s*2_ to *X*_*s*1_ = 0, instantaneous interaction = 0.463. **C**) The spectral decomposition of stochastic interaction, integrated information and Granger causality can be higher or lower than each other, depending on frequency (the green, blue and red lines can either above or below each other, depending on frequency). **D**) System with bidirectional and instantaneous influences. **E**) The measures in the time-domain. Stochastic interaction = 0.255, predictive information = 0.267, integrated information = 0.205, Granger causality from *X*_*s*1_ to **x**_*s*2_ = 0.086, Granger causality from *X*_*s*2_ to **x**_*s*1_ = 0.096, instantaneous interaction = 0.073. **F**) The spectral decomposition of the measures can all be higher or lower than each other, depending on frequency (the green, blue, red, maroon and yellow lines can all be above or below each other, depending on frequency). For the spectral decomposition of predictive information for these two systems see supplementary Figure S1C and D.

In the time-domain the strong instantaneous influence manifests as high instantaneous interaction, which leads to increased stochastic interaction (Eq. 37), such that the latter is now greater than predictive information (Figure 4B). In the absence of an influence from *X*_*s*1_ to **x**_*s*2_ Granger causality in that direction remains at zero. Notably, the strong instantaneous influence results in a higher value of integrated information than Granger causality from *X*_*s*1_ to **x**_*s*2_ (Figure 4B), whereas the two were previously identical (blue and red bars in Figure 3E are equal). This is counter-intuitive since the system has only a unidirectional causal influence and so total causal influence, as quantified by integrated information, should equal Granger causality from *X*_*s*1_ to **x**_*s*2_. In the supplementary information we demonstrate that we can better understand what is driving this difference by considering the structure of the transfer functions of the disconnected models (Figure S2).

The spectral decompositions of the measures display a complex interplay. Unlike the time-domain, stochastic interaction, integrated information and Granger causality can all be higher or lower than each other, depending on frequency (Figure 4C). Due to the strong instantaneous influence, instantaneous interaction remains above integrated information and Granger causality from *X*_*s*1_ to **x**_*s*2_ at all frequencies.

Particularly striking is the difference in the spectral decompositions of integrated information and Granger causality from *X*_*s*1_ to **x**_*s*2_, since the system is only unidirectionally-connected. These observations clearly demonstrate that the time-domain relationships between the measures (Figure 2) do not hold on a frequency-by-frequency basis for this system. We show in the Supplementary information that these differences are explained by differences in the diagonal entries of the disconnected models’ respective transfer functions.

We re-emphasize that the theoretical framework ensures that the sum over frequencies of the spectral decompositions equals the time-domain measures (Eq. 18). There is no theoretical guarantee that the spectral decompositions follow the time-domain relationships on a *frequency-by-frequency* basis.

Finally, we investigated a bidirectionally-connected system with instantaneous influences (Figure 4D), reflecting the most general system structure:

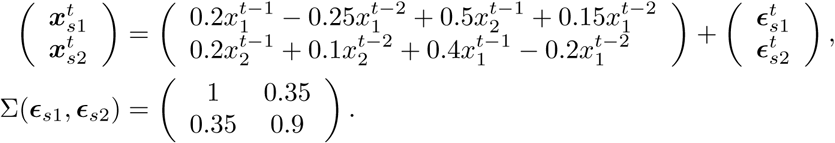

For this system, all the time-domain measures are non-zero and follow the time-domain relationships (Eqs. 36 and 37) (Figure 4E). The spectral decompositions once again display a complex interplay, in which any measure can be higher or lower than any other, depending on frequency (Figure 4F). This confirms that the spectral decompositions of the measures do not follow the time-domain relationships in this most general system structure.

## Discussion

In this paper, we presented a general framework for spectral decomposition of causal influences in linear autoregressive processes. The framework is based on a general information theoretic framework for quantifying causal influences in the time-domain [22] together with a result linking the generalized variance and the determinant of the spectral density matrices (Eq. 16), thus connecting the time-domain measures and their spectral decompositions. The spectral decompositions are such that the integral over frequencies equals the time-domain measures.

We used this framework to provide a spectral decomposition of integrated information, which has not been derived before. In addition, we used the framework to derive the spectral decompositions of Granger causality, stochastic interaction, predictive information, and instantaneous interaction in a unified manner. Spectral decompositions of Granger causality, total interdependence and instantaneous interaction have been previously described but our derivation here is novel. Thus, their relationship to our unified framework was not known *a priori*. In fact, we found that the spectral decomposition of instantaneous interaction we derived is different from a previous proposal [11] which lacked theoretical grounding [35]. We also confirmed that (1) the spectral decomposition of Granger causality we derived is the same as the one suggested by Geweke [7] and (2) that the spectral decomposition of stochastic interaction is the same as that of total interdependence [7, 11] and closely related to coherence. An important property of our derivation is that the spectral decompositions is applicable to any measure that can be defined by cutting a subset of the causal influences, making it directly applicable across arbitrary partitioning of the system.

In the time-domain, all the measures are non-negative because they are written as KL-divergence [22]. In addition, it has been shown that the spectral decomposition of Granger causality [7] and stochastic interaction [39, 40] are non-negative. We never observed the spectral decomposition of integrated information to be negative at any frequency across a variety of simulations. However, we observed that the spectral decomposition of predictive information we derived (Eq. 28) can become negative for some frequencies (Figure S1). Further theoretical work is required to understand the conditions under which the spectral decomposition is non-negative. However, the framework is sound in that the integral over frequencies equal the time domain measure. In addition, we demonstrated that the framework can be used to derive the well known spectral decomposition of Granger causality and show that the spectral decomposition of stochastic interaction is closely related to coherence. Thus, our framework unifies the derivation of quantities that have so far been considered in isolation. In sum, our proposed spectral decomposition of integrated information is theoretically solid and suitable for empirical investigation.

Using simulations, we showed that for a uni- and a bi-directionally connected system with instantaneous influences the spectral decompositions of the measure display a complex interplay in which every measures can be higher or lower than any other (Figure 4). We attributed this complex behavior to differences in the diagonal components of the respective disconnected models’ transfer functions (Figure S2). The complex frequency-by-frequency behavior of the spectral decompositions is contrasted with the time-domain, in which analytic relationships between the measures (Eqs. 36 and 37, Figure 2) guarantee an ordering of the measures such that the information loss increases with the extent of influences cut. Although our simulations of simpler systems show that time-domain relationships can hold on a frequency-by-frequency basis for specific systems (Figure 3), our more general simulations (Figure 4) show they cannot hold in general. We emphasize that our theoretical framework only guarantees that the integral over frequencies of the spectral decompositions equals the time-domain measures, so the observation that the spectral decompositions do not follow the time-domain measures on a frequency-by-frequency basis is theoretically consistent. While the relationships between the measures in the time-domain help in intuitively interpreting the measures, the departure from these relationships demonstrates that the spectral decompositions convey additional information about the system beyond that observable in the time-domain.

To conclude, the spectral decomposition of integrated information we derived provides a new tool for investigating frequency-specific causal influences. This will be particularly useful for systems for which frequency-specific causal influences are masked in the time-domain, as increasingly reported in neural systems [13,14,18,41,42]. In particular, the spectral decomposition of integrated information we presented offers a new angle for empirically investigating the integrated information theory of consciousness [43–49]. This will shed new light on how subjective conscious experience relates to the spectral characteristic of neural activity.

## Methods

### Computing the measures and their spectral decomposition

To calculate integrated information for a given autoregressive process, we first calculate the autocovariance function of the system (i.e. the autocovariance function of the full model) using the system’s autoregressive coefficient matrix *A* and residuals covariance matrix Σ(*ϵ*). This is efficiently computed using the var to autocov.m function from the Multivariate Granger Causality (MVGC) toolbox [34]. Next, the disconnected model’s autoregressive coefficient matrix *A*′ and residuals covariance matrix Σ(*ϵ* ′) are found by iterating through Eqs. 8 and 7 (under the constraint that *A*′ is block-diagonal). Integrated information is finally obtained by computing the log ratio of the generalized variances (Eq. 12). The spectral density matrix of the disconnected model *S*′(*ω*) is obtained following Eqs. 13 to 17. Once *S*′ (*ω*) is available, the spectral decomposition of integrated information is obtained by computing the log ratio of the determinants of the spectral density matrices (Eq. 20).

By imposing the relevant constraints on the disconnected models (Figure 1), the same procedure can be used to compute Granger causality and predictive information. In practice, however these measures can be calculated more efficiently. Efficient algorithms are available for calculating Granger causality, as implemented in the MVGC toolbox. Predictive information (Eqs. 27 and 28) can be calculated directly from the parameters of the full model. We have confirmed in simulations (not shown) that the different ways of computing these quantities give identical results.

Instantenous interaction (Eqs. 31 and 32) is calculated directly from the parameters of the full model. Time-domain stochastic interaction can be calculated once Granger causality has been estimated in both directions (Eq. 34). The spectral decomposition of stochastic interaction can be calculated directly from the spectral density matrix of the full model (Eq. 35).

The autoregressive order of the disconnected model is an important consideration in the estimation of integrated information. The disconnected model is fitted using the autocovariance function of the full model, so that the order needs to be chosen high enough so as to provide the best approximation of the full model. Failure to set the order sufficiently high could result in negative values and the violation of the time domain relations (Eqs. 36). Estimating the disconnected model parameters directly from the full model is also the approach taken in modern Granger causality analysis [34, 50, 51]. In our simulations we set the order of the disconnected models to the order of the autocovariance function (as computed by the MVGC toolbox). There was no discernible difference when setting the order any higher, confirming that this choice of order is sufficiently high.

Code for calculating the measures and running the simulations is available at https://github.com/drorcohengithub/PhiSpectralDecomposition.

## Supplementary Information

### The spectral decomposition of predictive information

**Figure S1.**
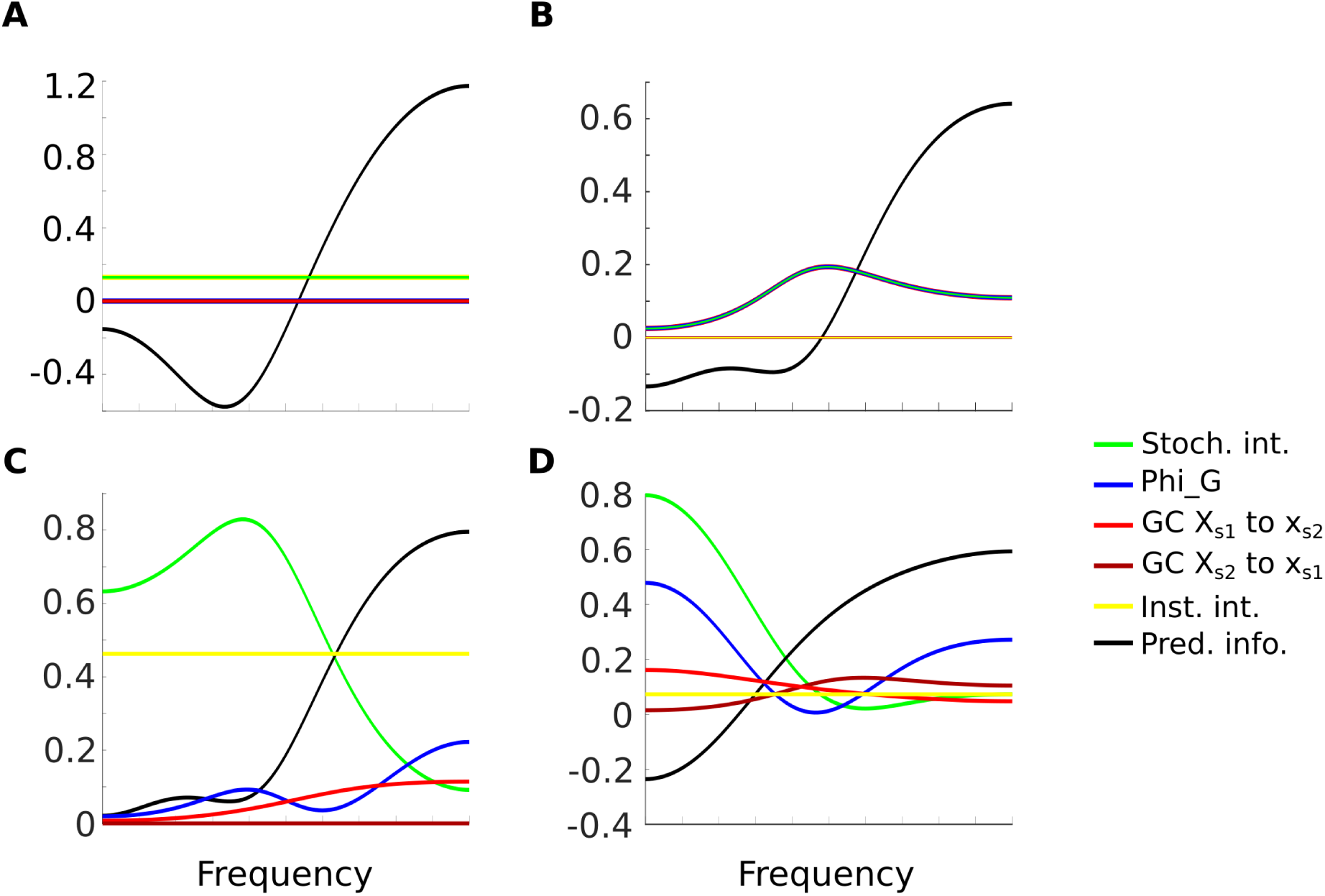
The spectral decomposition of predictive information (black) is negative for some systems and frequencies. The spectral decompositions of all other measures are also shown for comparison. **A**) The system with only instantaneous influence (Figure 3C). **B**) The system with only a unidirectional-influence (Figure 3F). **C**) The system with both a unidirectional and instantaneous influence (Figure 4C). **D**) The bidirectional system with instantaneous influence (Figure 4F).

### Explaining the differences between the spectral decompositions of the measures based on the transfer function

In our simulations we observed that the spectral decompositions of the measures display a complex interplay in which integrated information, Granger causality and stochastic interaction can be higher or lower than each other, depending on frequency. To better understand what is driving these differences, we rewrite the spectral density matrix for the disconnected model in terms of the transfer function

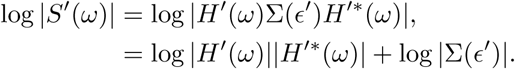

Since the transfer functions of the disconnected models used for stochastic interaction and integrated information are diagonal (Figure 1F and R), and the transfer functions for Granger causality is upper (or lower) triangular (Figure 1I), we can use that 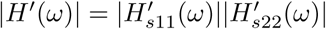. Thus we have

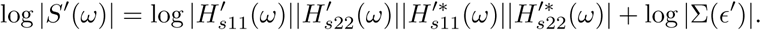

Focusing on bivariate systems as in our simulations we write

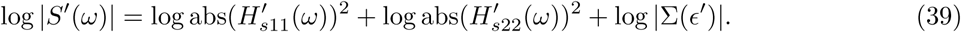

Substituting into the general spectral decomposition we obtain

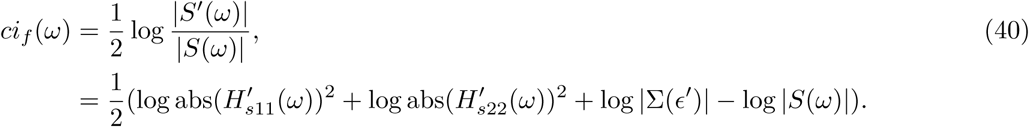

The log |*S*(*ω*) | term is identical for the spectral decompositions of all the measures. The term log |Σ(*ϵ*′)| is constant across frequencies and does not contribute to frequency-by-frequency differences. Thus, the frequency-by-frequency differences between the spectral decompositions are due to differences in the diagonal elements of the transfer function 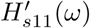 and 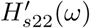.

We can use Eq. 40 to better isolate what is driving the differences between the spectral decompositions of the measures. For example, for the unidirectionally-connected system with instantaneous interaction (Figure 4A-C), there are two non-zero self-influences and one non-zero causal influence (Figure S2A), with corresponding non-zero components of the transfer function *H*_*s*11_(*ω*), *H*_*s*22_(*ω*) and *H*_*s*12_(*ω*). Each of these components has a particular spectral profile (Figure S2B). Since this system is unidirectionally-connected (i.e. the transfer function is upper triangular), we can apply the decomposition in Eq. 39 to the original spectral density matrix

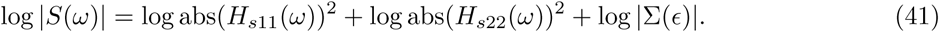

The term log *|*Σ(ϵ)*|* represents an offset across all frequencies and does not affect frequency-by-frequency behaviour (Figure S2C). Finally, from Eq. 41 we can construct log *|S*(*ω*)*|* (Figure S2D).

The structure of the disconnected model used for quantifying Granger causality from *X*_*s*1_ to **x**_*s*2_ contains an influence from *X*_*s*2_ to **x**_*s*1_, but no influence in the opposite direction (Figure S2E). As a result, 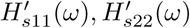 and 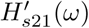 are non-zero, whereas 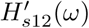 is forced to zero (Figure S2F). Note that even though there is no influence from *X*_*s*2_ to **x**_*s*1_ in the original system, this influence is present in the disconnected model. This observation reflects that the best possible approximation for the original system in the time-domain (i.e. across all frequencies) under the constraint that there is no influence from *X*_*s*1_ to **x**_*s*2_, may include an influence from *X*_*s*2_ to **x**_*s*1_. Since the disconnected model is only an approximation, we have that log |Σ(*ϵ*) |*<* log Σ(*ϵ*′) | (Figure S2G). From Eq. 39 we can construct log *|S*′ (*ω*)*|* (Figure S2H). Finally, Figure S2I shows (half of, Eq. 20) the difference between log *|S*(*ω*)*|* and log *|S*′ (*ω*)*|*, which reflects the difference of the terms 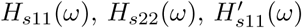 and 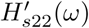 in Eqs. 39 and 41. For this disconnected model the subtraction reflects the spectral decomposition of Granger causality from *X*_*s*1_ to **x**_*s*2_ (Figure S2I, identical to red line in Figure 4C).

The disconnected model used for integrated information does not have an influence from *X*_*s*1_ to **x**_*s*2_ or from *X*_*s*2_ to **x**_*s*1_ (Figure S2J). Thus, the only non-zero components of the transfer function are 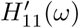 and 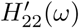 (Figure S2K). The structure of the transfer function used to estimate integrated information is different from that used for Granger causality, since the latter also included a non-zero influence from *X*_*s*2_ to **x**_*s*1_ (Figure S2F). The offset due to log | Σ(*∊*′) | is greater again, as per the time domain relations between the measures (Eq. 36) (Figure S2L). As before, the differences in 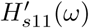 and 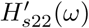 between the original and disconnected model lead to a difference in log |*S*(′*ω*)| (Figure S2M), resulting in a different spectral profile for integrated information (Figure S2N, identical to blue line in Figure 4C).

The structure of the disconnected model for stochastic interaction does not allow any causal or instantaneous influences (Figure S2O). The non-zero components of the transfer function are 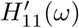 and 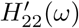 (Figure S2P). Though these are also the only non-zero components of the transfer function used for integrated information, their spectral profiles are not the same (compare Figure S2K with P). This demonstrates again that the best possible approximation for the original system in the time-domain under the constraint that there are neither causal nor instantaneous influences (stochastic interaction) leads to a different transfer function than that under the lesser constraint of only no causal influences (integrated information). For stochastic interaction, we observe a much greater value of log *|*Σ(*ϵ*′) *|*, which is attributed to the strong instantaneous interaction in this system (Figure 4A and Eq. 37) (Figure S2Q). In turn, this leads to a generally greater magnitude for log |*S*′(*w*)| as compared to that of Granger causality and integrated information (Figure S2R). In high frequencies however, the negative values of log 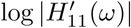 and log 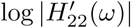 (frequencies below dotted line in Figure S2P) work against the high value of log |Σ(*ϵ*′)|, such that stochastic interaction in this region is in fact smaller than integrated information (Figure S2S).

This analysis demonstrates how differences between the spectral decompositions of the measures can be explained in terms of differences in the diagonal components of the transfer function 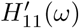 and 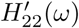. The best approximations for the original system in the time-domain have different transfer functions, strongly depending on the specifics of the constraints applied. One complicating factor is that the terms 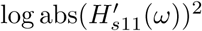 and 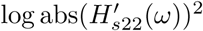 can have different signs at different frequencies, such that they can either compound or cancel. This can make it challenging to quantify their individual contribution to the spectral decomposition of the measures.

**Figure S2.**
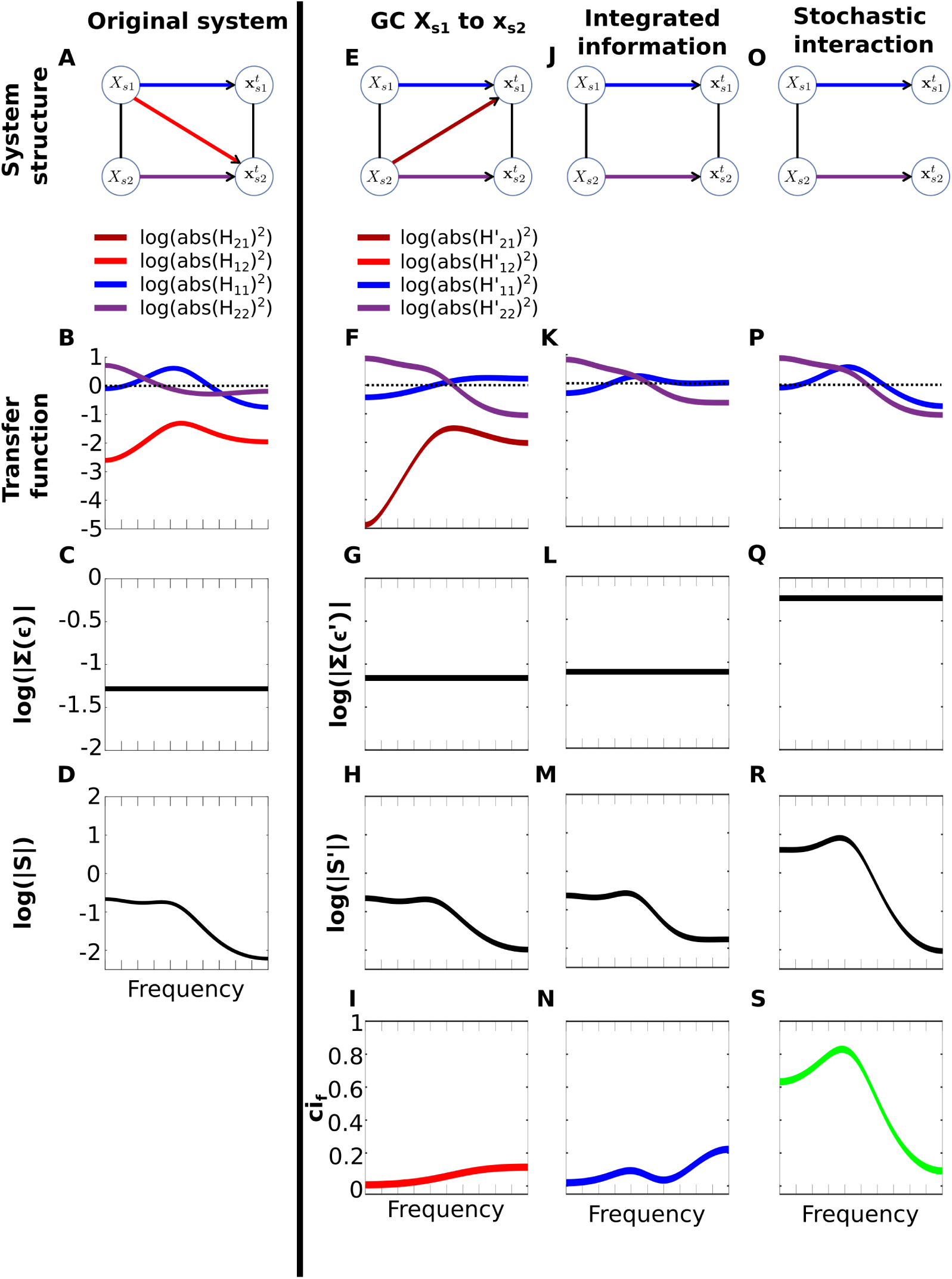
The differences between the spectral decompositions of the measures can be attributed to differences in the structures of the measures’ respective transfer functions. The figure represents the decomposition in Eq. 40 applied to the system in Figure 4A-C (see text for details). **A**) Structure of unidirectionally-connected system with instantaneous interaction. Same as Figure 4A but with color-coded self- and causal-influences. **B**) The spectral profile of the components of the transfer function (see legend, the frequency dependence *ω* has been omitted). Since the system is unidirectional *H*_12_ is not shown. The dotted line represents *y* = 0. **C**) log *|*Σ(ϵ)*|*. **D**) log *|S*(*ω*)*|* obtained following the spectral decomposition in Eq. 40. **E** – **H**) same as (**A** – **D**) for the disconnected model used to compute Granger causality from *X*_*s*1_ to **x**_*s*2_. **H**) log *|S* ′ (*ω*)*|*.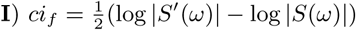, providing the spectral decomposition of Granger causality. (**J – N**) and (**O – S**) same as (**E – I**) for the disconnected models used to compute integrated information and stochastic interaction, respectively.

### Constraints for the disconnected model

When we cut the causal influence from the element *n* to the element *m*, we impose the following constraint [22],

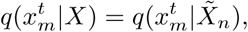

where 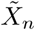 is the complement of *X*_*n*_ in the whole system *X*. This constraint means that there is no direct influence from the node *n* to node *m* given the states of the other elements 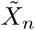 being fixed. 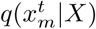 is determined by the autoregressive model,

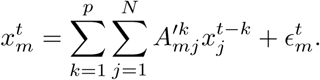

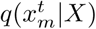 is explicitly given by

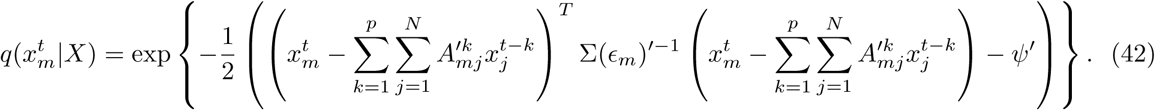

When 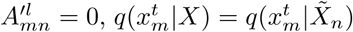 because *X*_*n*_ does not appear in the right hand side of Eq 42. Also, 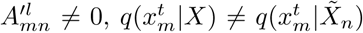 because *X*_*n*_ does appear in the right hand side of Eq 42. Thus, the constraint 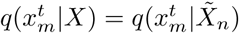 is equivalent to

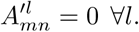

### Derivation of instantaneous interaction

Since the constraint for instantaneous interaction (Eq. 29) was not examined in [22], we derive the minimized KL here. Under this constraint, the KL divergence between the full and disconnected model simplifies to

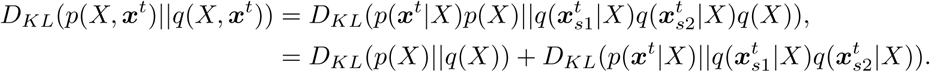

The first term vanishes when *q*(*X*) = *p*(*X*). Expanding the second term, we obtain

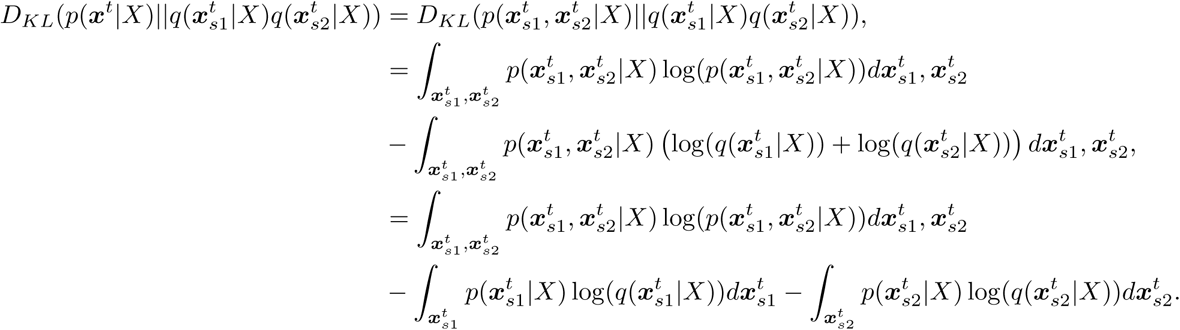

The first term is the entropy of *p*. It does not depend on *q* and can be ignored when minimizing the above KL divergence. We can rewrite the remaining two terms as the sum of KL divergences and entropies

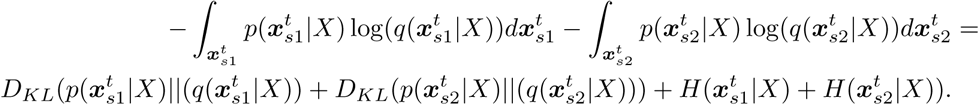

Since the entropy terms are independent of *q*, this expression is minimized when 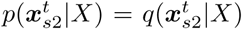 and 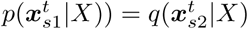. The minimized KL divergence is

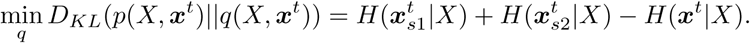

Note that (1) the factorization 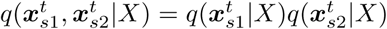 implies that Σ(*ϵ*′) is diagonal and (2) that 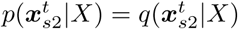 and 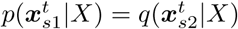 are satisfied when *A* ′ = *A*.

### Equivalence to spectral Granger causality

Our goal is to show that when using the disconnected model for Granger causality (from *X*_*s*1_ to 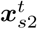)

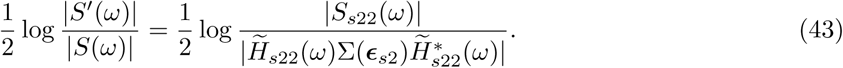

To prove this we will show that there is an analytic relationship between the entries of the transfer function of the full model *H* and the transfer function of the disconnected model *H*′. The relationship is such that the lhs simplifies to the rhs. Before establishing this relationship we first rewrite the lhs in terms of the transfer functions (omitting the frequency dependence *ω*)

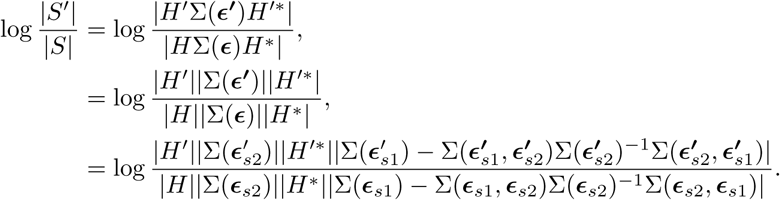

where for the last line we applied the formula for the determinant of a block matrix to *|*Σ(*ϵ*)*|* and *|*Σ(***ϵ*′**)*|*,

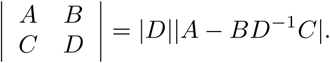

Since *H*′ is upper triangular, we can use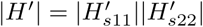. Combined with 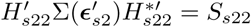 we have

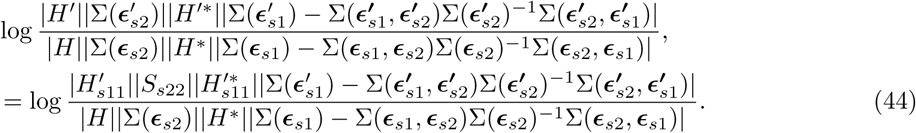

To link with the transformation from *H* to 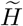 in the definition of Granger causality (rhs of Eq. 43), we consider the matrix *P*

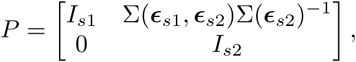

where *I*_*s*1_ and *I*_*s*2_ are appropriately sized identity matrices and 0 is an appropriately sized zero matrix. *P* is such that

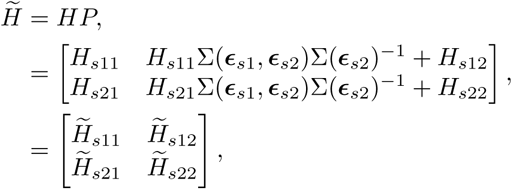

where 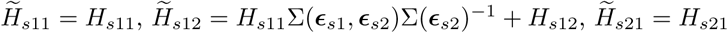 and 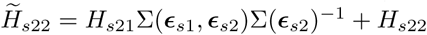

Since *|P |* = 1, we can substitute 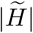 for *|H|*. Using this substitution together with applying the formula for the determinant of a block matrix to 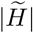 we can rewrite Eq. 44 as

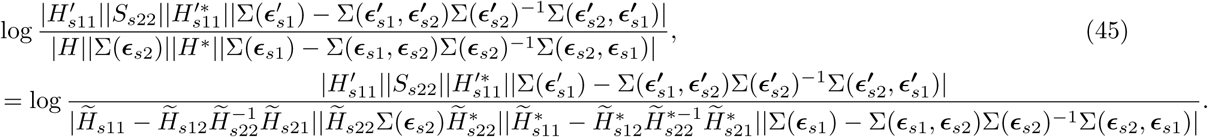

If we can show

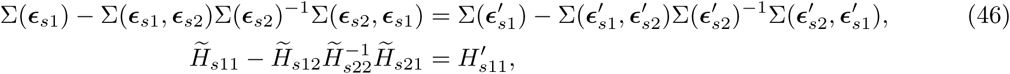

then Eq. 45 simplifies to log 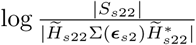, completing the proof.

To show these, we first recall that minimizing the KL between the full and disconnected model gives [22]

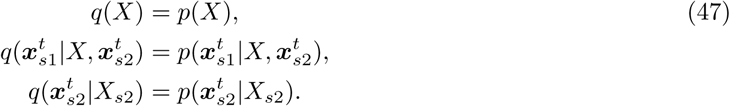

For zero-mean Gaussian systems [8]

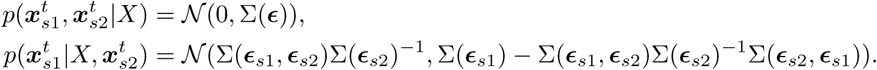

Using that 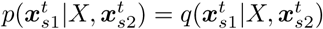 (Eq. 47), we find

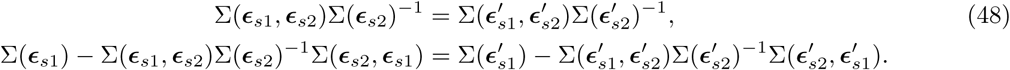

This proves the first line of Eq. 46.

To prove the second line of Eq. 46 we examine what the equivalence in Eq. 48 means for the autoregressive coefficients of the models. We begin by considering the transformation of the residuals by *P* ^*-*1^Σ(*ϵ*)(*P* ^*-*1^)^*T*^

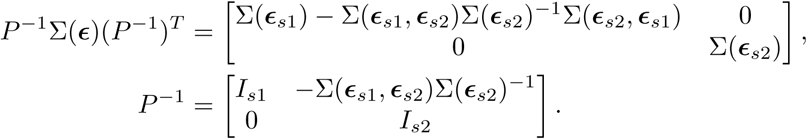

Since 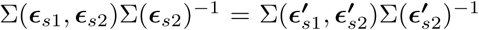 (Eq. 48), we have *P* = *P*′. Thus, for the disconnected model

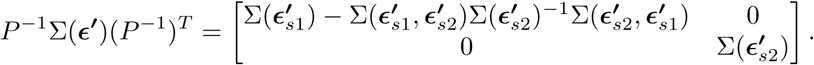

We can see that the top left entries of *P* ^*-*1^Σ(*ϵ*)(*P* ^*-*1^)^*T*^ and *P* ^*-*1^Σ(*ϵ* **′**)(*P* ^*-*1^)^*T*^ are equivalent (Eq. 48). The difference between the transformed residuals is

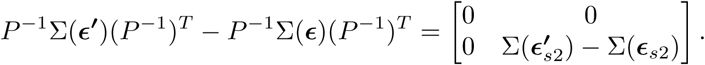

Next we express this in terms of the autoregressive coefficients. We have shown that (Eq. 10)

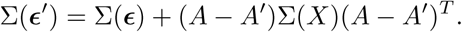

Re-arranging and applying the transformation we have

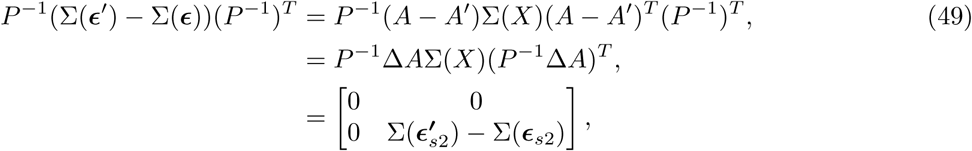

where we used

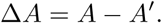

Equating the top left entries of Eq. 49 we find

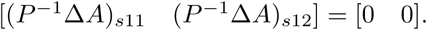

Using (*P* ^*-*1^Δ*A*)_*s*11_ = 0 and from the definition of the transfer functions (Eq. 15) we have in the frequency domain

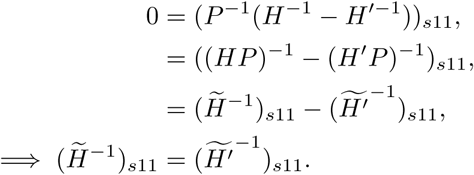

In terms of the inverse of 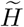

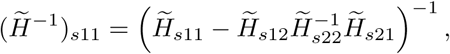

which follows from the inverse of a block matrix

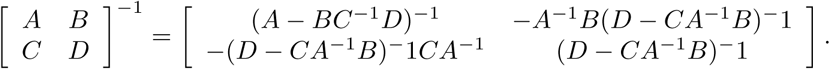

In terms of the inverse of 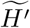

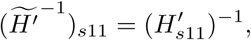

which follows since 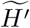 is upper triangular.

Putting these together we have

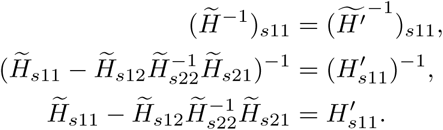

proving the second line of Eq. 46, as required.

